# A robot-based gait training therapy for pediatric population with Cerebral Palsy: goal setting, proposal and preliminary clinical implementation

**DOI:** 10.1101/255448

**Authors:** C. Bayón, T. Martín-Lorenzo, B. Moral-Saiz, O. Ramírez, A. Pérez-Somarriba, S. Lerma-Lara, I. Martínez, E. Rocon

**Author notes:** Corresponding author: Eduardo Rocon, Centro de Automática y Robótica, Consejo Superior de Investigaciones Científicas, Ctra Campo Real km 0.2 - La Poveda-Arganda del Rey, 28500 Madrid, Spain.

## Abstract

**BACKGROUND:** The use of robotic trainers has increased with the aim of improving gait function in patients with limitations. Nevertheless, there is an absence of studies that deeply describe detailed guidelines of how to correctly implement robot-based treatments for gait rehabilitation. This contribution proposes an accurate robot-based training program for gait rehabilitation of pediatric population with Cerebral Palsy (CP).

**METHODS:** The program is focused on the achievement of some specifications defined by the International Classification of Functioning, Disability and Health framework, Children and Youth version (ICF-CY). It is framed on 16 non-consecutive sessions where motor control, strength and power exercises of lower limbs are performed in parallel with a postural control strategy. A clinical evaluation with four pediatric patients with CP using the CPWalker robotic platform is presented.

**RESULTS:** The preliminary evaluation with patients with CP shows improvements in several aspects as strength (74.03±40.20%), mean velocity (21.46±33.79%), step length (17.95±20.45%) or gait performance (e.g. 18.88±14.31% in Gross Motor Function Measure-88 items, E and D dimensions).

**CONCLUSIONS:** The improvements achieved in the short term show the importance of working strength and power functions meanwhile over-ground training with postural control. This research could serve as preliminary support for future clinical implementations in any robotic device.

**TRIAL REGISTRATION:** The study was carried out with the number R-0032/12 from Local Ethical Committee of the Hospital Infantil Niño Jesús. Public trial registration: ISRCTN18254257. Registered 23 March 2017, retrospectively registered.

## Introduction

Gait limitation is one of the main impairments in children with Cerebral Palsy (CP) [1]. This mobility deficiency in CP is commonly the consequence of a damage of the child’s Central Nervous System (CNS), and an optimal functional training is required in order to maximize the improvements [2], which will highly contribute to the enhancement of the independence and, therefore, the quality of life for both the young patient and his family [3].

The use of robotic trainers for neurorehabilitation applications has increased in the last decades, both in adulthood and childhood, and in several motor diseases [3–5]. Robot-based therapies have been developed and improved beyond reducing the clinician’s effort. Currently, a new generation of robotic devices [6–8] provides means for encouraging the patients to an active participation in exercises, which are now more task specific. Both the implemented novel control strategies and the modularity of new exoskeletons and gait trainers offer promising possibilities to enhance the rehabilitation outcomes by adapting the treatment to the patient’s needs [9,10]. Nevertheless, so far there is not enough evidence to ensure that classic robot-based rehabilitation provides better treatment outcomes by itself than conventional physical strategies in childhood [11]. New approaches are needed in order to improve the rehabilitation, making the robotic therapy a key feature of the change.

One of the main drawbacks for the everyday use of these technologies into the rehabilitation practice, apart from the price of these devices, is the absence of studies that describe a detailed robotic training program for gait rehabilitation. The wide variety of changes that could be applied to the parameters of robotic training therapies, makes unclear how to specify rehabilitation settings with the aim of providing a suitable solution for a large population size. Additionally, most of current studies are only focused on lower limbs strategies. However, the upper body (head and trunk movement) also influences gait function through walking balance [12], so a proper program should not ignore these features.

This manuscript presents a detailed robot-based therapy proposal for the rehabilitation of gait function in children with CP, which is based on the achievement of some specifications defined by the International Classification of Functioning, Disability and Health framework, Children and Youth version (ICF-CY) [13]. It contributes with better answers on how to implement robotic rehabilitation following defined guidance, establishing the baseline settings and subsequently tailoring the therapy to each patient.

The proposed robotic rehabilitation therapy works around a key factor: the implementation of strength and power exercises at the same time than over-ground walking guidance, performing in parallel an active head-trunk control therapy. As a result, the robot-based program recreates a situation as similar as possible to a real gait scenario, and encourages the patients to control different movements associated with gait: not only individual movements of lower limb joints but also the synergy between them while maintaining a proper posture of upper body. We hypothesize that these essential components, performed following an appropriate progression of the variables, will boost the rehabilitation of our patients.

Eventually, the robot-based therapy proposed in this manuscript is evaluated in four patients with CP in order to provide preliminary results of its application. Although the proposed robot-based treatment could be implemented in any robotic device adapted for that, in this study authors selected the CPWalker training platform [7] as one scenario to test the effectiveness of the approach. This device is a novel robotic prototype with partial body weight support (PBWS) for over-ground rehabilitation of children with CP. It provides means for adapting the therapy to the user’s necessities through different levels of assistance in multi-joint over-ground training. Other gait trainers, such as Lokomat [14] or LOPES [6], could have been also used to evaluate this training method, since they are also able to tailor each therapy session to the patient’s needs.

## Materials and Methods

### Rehabilitation device

The CPWalker rehabilitation platform [7] is a robotic device composed by an exoskeleton linked to a walker that provides support and balance to the child during over-ground training (Figure 1). The device is able to implement users’ PBWS and allows the adaptation of exercises to the patient’s capabilities by means of individual controllers for each joint, which increases the modularity of the system [7]. Each joint of the CPWalker exoskeleton can operate in a wide range of modes (see Figure 1):

i. *Position control mode*: in this mode, the robot imposes a prescribed gait pattern to the user’s lower limbs. The aim is that the patient learns the walking motion sequence correctly.
ii. *Impedance control modes*: these modes take into account the patients’ collaboration. Thereby, the prescribed gait pattern should be achieved by the sum of robotic assistance and patient’s cooperation. Three different modes of impedance may be executed in CPWalker (high, medium and low), which tolerate variable deviations from the programmed gait trajectories, enhancing the patients’ participation and taking advantage of their residual movements through assist as needed strategies (AAN).
iii. *Zero-force control mode*: in this mode, the trajectory reference is not given, and the patient is who entirely moves the legs with a minimal resistance of the exoskeleton. It is used with patients with enough motor control (acquired with the previous modes) but poor balance, so the CPWalker provides stability and PBWS while the patient implements the gait pattern.

**Fig. 1.**
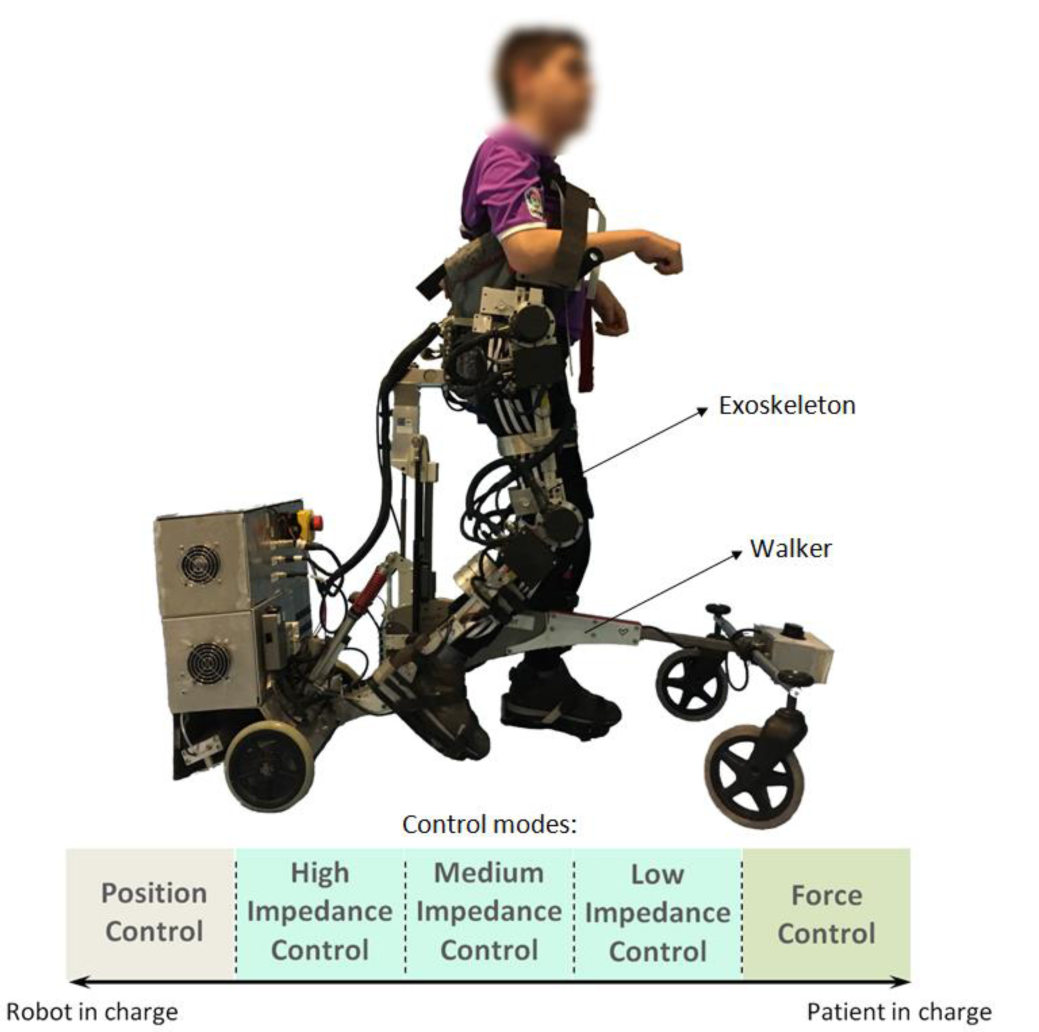
**CPWalker robotic platform (*exoskeleton* to guide the patients’ lower limbs and *walker* with PBWS to provide balance control during over-ground walking). The wide variety of operating modes comprehends from “robot in charge” to “patient in charge” states: i) position control mode; ii) high impedance control mode; iii) medium impedance control mode; iv) low impedance control mode; v) zero-force control mode**.

Within each mode, some variables of the robotic platform are updated along the treatment sessions: PBWS, gait velocity and the percentage of the total range of motion (ROM) [9]. The possibility of varying these parameters enables the customization of the therapy to the progression of each patient and gives a higher versatility for the treatment design. Additionally, the robotic platform also includes a biofeedback strategy to motivate the children to actively correct their posture during walking [7,9]. The CPWalker robotic platform may be easily controlled by a clinician through an intuitive interface, which controls and monitors the exercises in real time.

The variety of control modes, the number of adjustable parameters, the ease of setting the variables individually for each joint and the possibility of implementing different strategies associated with gait simultaneously, made CPWalker an appropriate platform for testing the robotic therapy proposed in this manuscript.

### Robotic training program

In order to define the objectives of the robot-based treatment, the authors adopted the conceptual framework of ICF-CY [13]. The proposal was focused on improving the principal gait-related functions derived from this international classification. Concretely, the selected goals of the ICF-CY to be achieved with the treatment and the work methodology implemented in the robotic device (in this case CPWalker), are presented in Table 1.

**Table 1.** **Goal settings of the ICF-CY and the robot-based solutions adopted with CPWalker platform**

| ICF-CY<br>functional domain<br>[13] | Implementation on CPWalker |
| --- | --- |
| Mobility and stability of joint functions (b710, b715) | The different control modes of the exoskeleton are used to guide the movement of a single or multiple joints, improving motor control. The exoskeleton also helps to maintain stability through the coordinated actions of surrounding tissues. This domain is exercised along the whole treatment with diverse robotic assistance. |
| Muscle power functions (b730) | The *second phase* of the training requires the patient to contract a muscle or muscle groups to generate the necessary force in order to start and maintain the movement with AAN strategies. The force must be maintained for a time in the extremes of the gait pattern (maximum flexion and extension) in order to reach the these maximum values in the complete range of motion. |
| Muscle endurance functions (b740) | Muscle endurance is exercised when the patient is requested to sustain a muscle contraction to finalize the required movement with AAN strategies, mainly in the extremes of the gait pattern (maximum flexion and extension). |
| Control of voluntary movement functions (b760) | The voluntary movement is implemented through the control and coordination of simple and complex movements to collaborate with AAN strategies. Lower impedance implies more patient's control. |
| Gait pattern functions (b770) | Motor control and gait pattern functions are trained through the different control modes to guide the lower limbs following prescribed gait patterns at |
|  | several velocities and supports. |
| Maintaining a body position (d415) | A biofeedback strategy for postural control is used to notify the patients when they lose the correct position of the upper body. See section "Postural control". |
| Walking (d450) | Over-ground walking training is executed in all sessions with controlled PBWS, at different velocities and supports. |
| Proprioceptive functions (b260) | During random moments of the first training session, the patients perceive feeling using a mask on the eyes at the same time than the robot performs the movement for single or multiple joints with 100% of PBWS. |
| Formal relationships (d740) | Creating and maintaining patient-researcher relationship. See section "Motivation and inclusion of challenges". |

To achieve the goals presented in Table 1, authors previously performed a systematic selection of variables based on the requirements of the National Strength and Conditioning Association (NSCA) youth training guidelines, which suggests that eccentric and explosive strength exercises should be the beginning of a proper training to ensure greater muscle power generation and the transference of gains to gait (Table 2).

**Table 2.** **National Strength and Conditioning Association (NSCA) youth training guidelines** [15]

| ADAPTED YOUTH RESISTANCE TRAINING NSCA GUIDELINES |  |  |
| --- | --- | --- |
| Variables | Strength | Power |
| Muscle actions | Eccentric / Concentric | Eccentric / Concentric |
| Exercise | Single / Multi-joint | Single / Multi-joint |
| Intensity | ↑ Load / ↓ Velocity | ↓ Load / ↑ Velocity |
| Velocity | Moderate | Moderate / Fast |

According to the proposed objectives (Table 1) and complying with the NSCA youth training guidelines (Table 2), the treatment was conceptualized into two main *phases*, where the ROM, PBWS and gait velocity were the principal parameters under variation. The intention was that the patient maximized the gains acquired in the whole rehabilitation period (ideally the sum of robot-based exercises and common non-robotic therapy). A detailed description of each *phase* follows:

i. *First phase*: the main aim of this *phase* was to improve motor control, teaching the patients the correct sequence of motion and increasing strength. The patients were requested to follow the movements established by the exoskeleton with the minimal possible resistance during swing period, pushing the ground at each step and trying to keep the maximum flexion-extension values at the end of each gait cycle. Instructions were given to ensure the comprehension of normative gait patterns, and verbal encouragement in addition to direct feedback by graphics in real-time was delivered throughout the sessions.
ii. *Second phase*: the main aim of this *phase* was to further train motor control and increase power in order to ensure the transference to the independent gait pattern. Aware of the sequence of movement of a normal gait pattern, the patient’s contribution became an important aspect to develop neuroplasticity and preserve the gained motor control [16]. The active participation was achieved both by boosting the patient’s motivation and by requiring self-activity [17]. The latter was implemented through AAN algorithms and the impedance control modes presented previously for CPWalker [7].

Figure 2 represents the schematic view of the therapy proposal. The treatment was composed of a total of 16 sessions (*first phase*: 8 sessions for strength training and motor control learning; and *second phase*: 8 sessions to transfer the gains to gait through power performance). Exercises were multi-joint and gait-oriented, demanding concentric-eccentric actions based on the gait phase that was being performed. During the whole treatment (*first* and *second phases*) the position of head and trunk during walking exercises were monitored, especially because these patients usually walked looking at the ground. For this purpose, the strategy for postural control of CPWalker encouraged the patients giving an acoustic feedback to when their position was inappropriate, so they could realize and rectified it by themselves. The program was reinforced with modifications on AAN levels according to the patient’s evolution based on performance evaluations related to ROM, PBWS and gait velocity. Furthermore, several challenges were included to enhance the patients’ motivation.

**Fig. 2.**
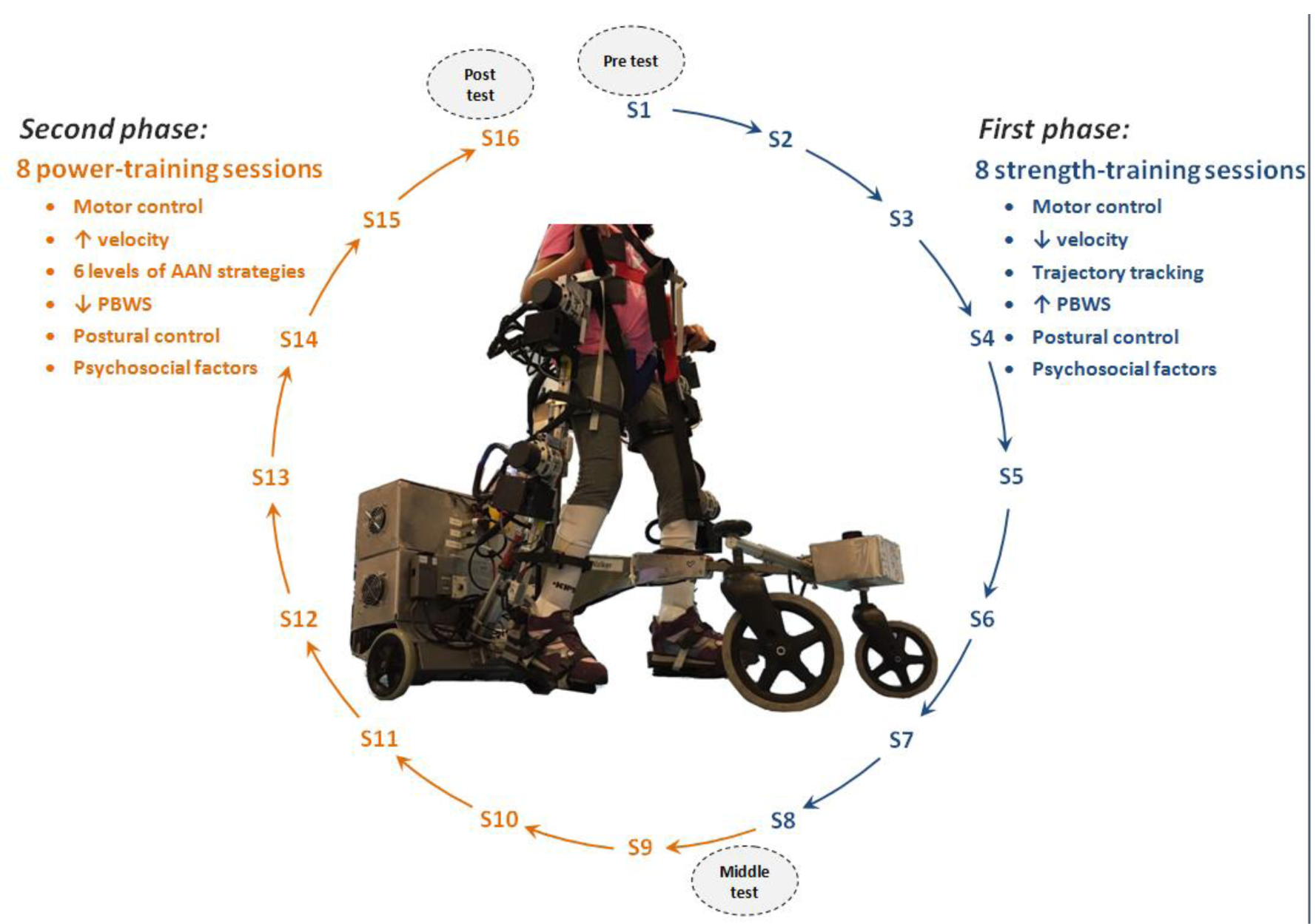
**Robot-based training program overview. *First phase:* sessions S1 to S8 for strength exercises where motor control was primary trained. *Second phase:* sessions S9 to S16 for power training where the assistance was progressively decreased with the patient’s progression. The improvements were assessed at three stages of analysis: before treatment begins, between both *phases* and at the end of the program (gray ellipses in the figure)**.

#### Duration of the study

The robot-aided treatment was proposed for a whole period of 2 monthly cycles (one month for each *phase* of the treatment) with the aim of having enough sessions to generate significant neural changes [18]. The children trained 2 non-consecutive days per week for 8 weeks (16 sessions, see Figure 2). The sessions consisted of a 10-15 minutes warm-up and 60 minutes of over-ground exercise with CPWalker, including 3 minutes of independent gait as a cool-down phase.

As Figure 2 indicates, the first 8 sessions corresponded with general motor control and strength exercises, where the robot imposed a gait trajectory tracking. Sessions 9 to 16 were related to muscle power performance through levels of AAN strategies, where self-activity was required.

#### Training phases

In order to individually define the training progression through the different sessions and to comply with the NSCA guidelines, the principal modifications were implemented on ROM, PBWS and gait velocity. The selected parameters for both *phases* are represented by Figure 3 and Figure 4 respectively. This selection was concluded in collaboration with our clinical partners, based on the evaluation of previous studies carried out with CPWalker [9,19]. The robot-based tasks began with high assistance and PBWS, and they progressed toward greater ROM and smaller PBWS as long as the patient overcame the different levels of the sessions. Parameter variations within and between sessions were performed as long as session goals were attained and when the clinical staff agreed, based on levels of spasticity, fatigue and motor control presented in the last day. If the patient was not ready to jump to the next challenge, the session was repeated with the same last percentages.

**Fig. 3.**
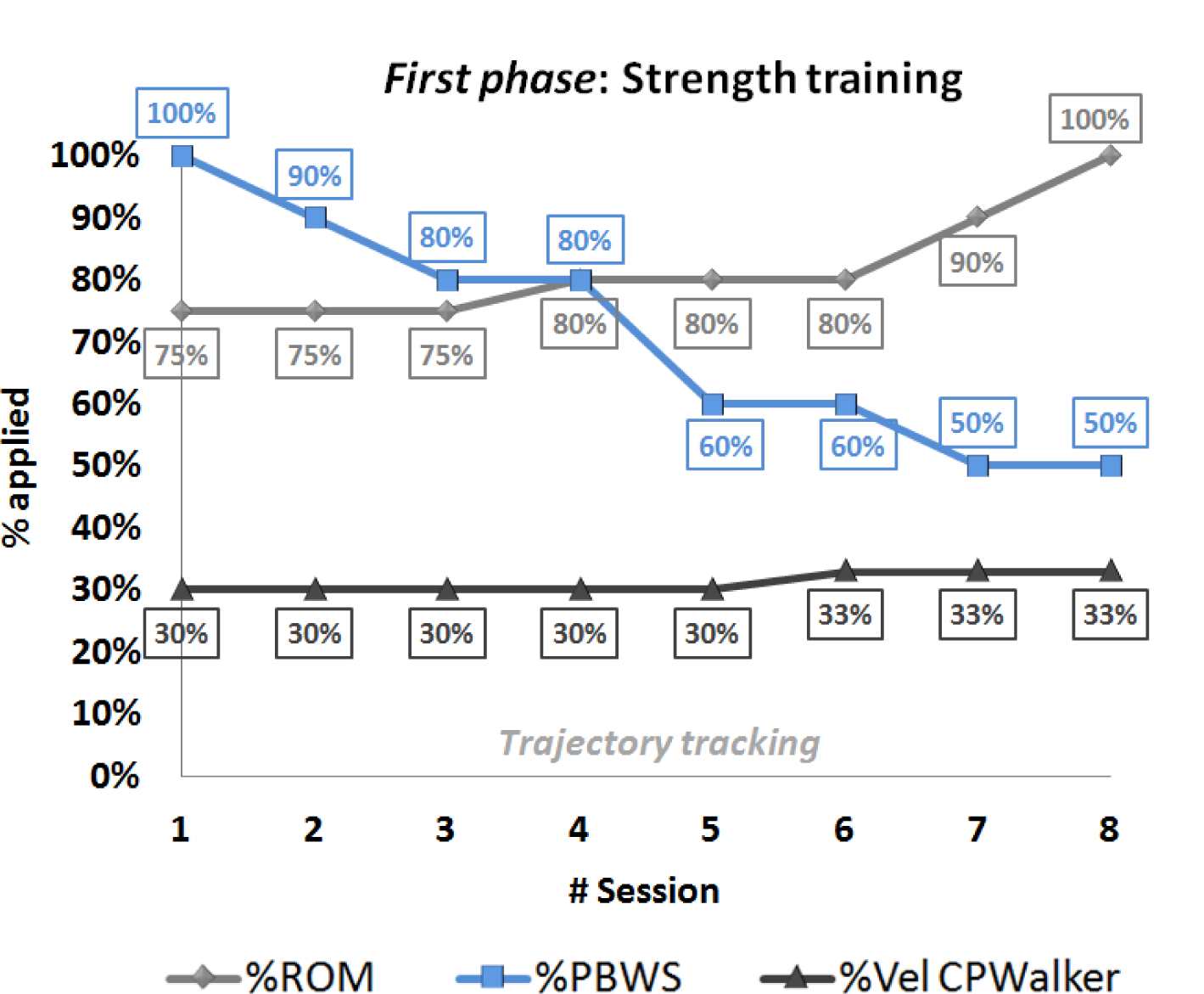
***First phase*: strength training progression values along first 8 sessions of the robot-based therapy. Trajectory tracking motion was imposed by the robot. The light-gray line represents the movement amplitude (%ROM), the blue line the changes of %PBWS and the dark-gray line is referred to gait velocity percentage for each session**.

**Fig. 4.**
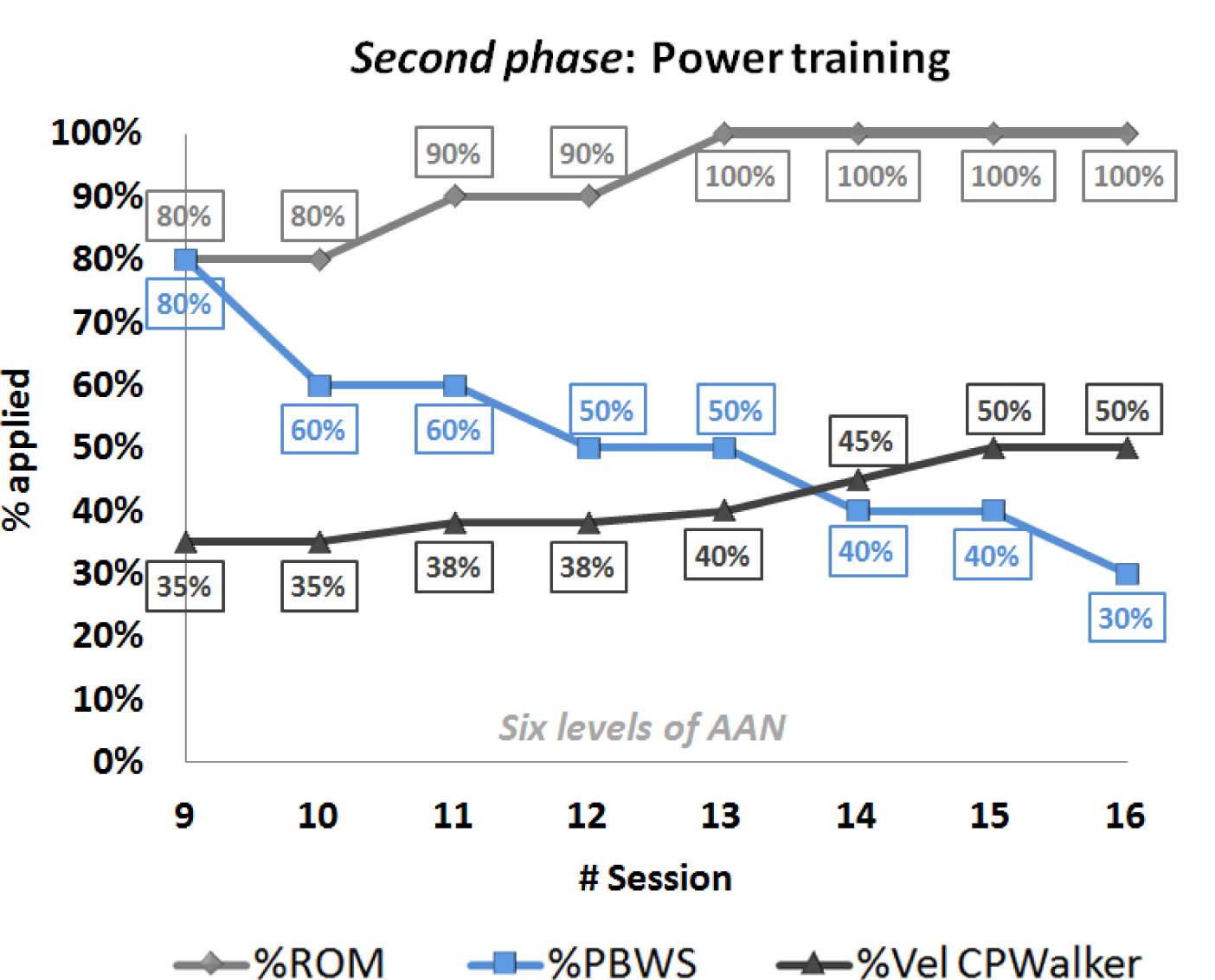
***Second phase*: power training progression values along sessions 9 to 16 of the robot-based therapy. Six different levels of assistance were selected on the exoskeleton (combining hips and knees). The light-gray line represents the movement amplitude (%ROM), the blue line the changes of %PBWS and the dark-gray line is referred to gait velocity percentage for each session**.

Within the *first phase* (Figure 3), the first session was performed with the children completely suspended (100% of PBWS) in order to adapt the users to the movements with the robot. Moreover, during random moments of this first session, the patients wore a mask on their eyes to feel the motion performance. For the rest of the *first phase* children’s lower limbs were guided through a pure position control imposed by the CPWalker platform, with a gradual decrease in PBWS (Figure 3 blue), and a gradual amplification of ROM (Figure 3 light-grey). Gait velocity during this *first phase* was maintained around a regular and small value (Figure 3 dark-grey). Notice that in general, only one variable at each session was varied.

The *second phase* of the training (sessions 9 to 16 in Figure 4) presented an additional difficulty that enhanced the user’s collaboration in the exercises performance through different levels of AAN strategies in the exoskeleton. The initial ROM for the *second phase* was set at 80% of the total gait pattern and reached 100% by session 13 (Figure 4 light-grey), time at which velocity was highly increased (Figure 4 dark-grey). Note that gait velocity for this *phase* became around double of the one achieved in the *first phase*, which is in relation to the requisites exposed in Table 2. Furthermore, PBWS declined up to 30% of weight supported by the platform (Figure 4 blue).

It is important to highlight in Figure 3 and Figure 4 that the percentage of PBWS was related to the individual patient’s total weight, and the percentage of ROM was applied to the total trajectory of the gait pattern programmed in the control of CPWalker [7]. The estimated changes in gait velocity are represented regarding percentages of CPWalker platform, where 100% corresponded to 0.6 m/s.

### Tailored Assist as Needed strategies

With the aim of enhancing the patient’s participation in the *second phase* and consequently improving outcomes of the treatment, in sessions 9 to 16, position control was substituted by six adapted levels of impedance control in the joints of the exoskeleton (Table 3).

**Table 3.**
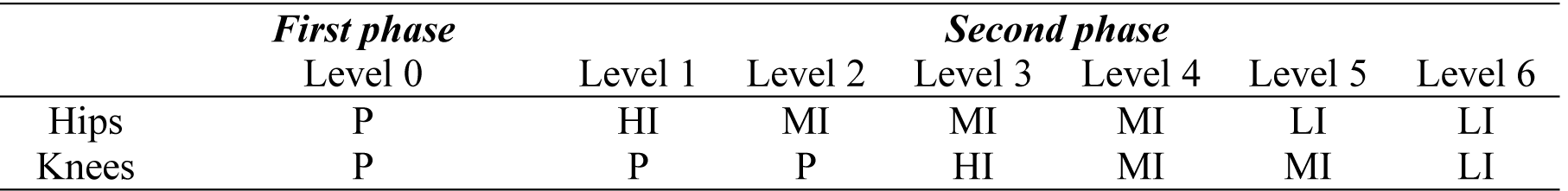
**Levels of assistance in *first* and *second* training *phases*: Position (P); High Impedance (HI); Medium Impedance (MI); Low Impedance (LI). The assistance on knee joint always went behind the assistance on hip, due to the knee movement during gait is performed following inertial forces, assigning to the hip movement higher importance. A higher level is implemented if the session performance (real motion versus desired pattern) is bigger than 85%**.

One of the main advantages of CPWalker robot was the possibility of selecting three different modes of assistance individualized per joint beyond a pure position control (high, medium and low impedances). In consonance with this, six situations (levels) were adopted following the criteria of our clinical partners in order to define the scales of difficulty in the assistance of knees and hips in the *second phase* (Table 3). Thereby, the patients were considered fit to move to the next level when they achieved a performance higher than 85% in the execution of each session, together with its corresponding parameters (ROM, PBWS and gait velocity) represented by Figure 4. This percentage of performance was calculated comparing the real motion executed by the children and the desired gait pattern of each session.

#### Postural control

It was important to ensure that throughout all the sessions, patients maintained a proper posture of head and trunk because it facilitates the performance of any activity of daily living, and improves the social interaction, the participation and communication [12,20]. In this sense, the aim of our proposal was to provide biofeedback to the patients each time that they kept an incorrect position of the body during walking. With this goal, the CPWalker robot used Inertial Measurement Units (IMUs) (Technaid, Spain) to measure the rotation of head and trunk in real time, and give acoustic feedback when subjects overcame predefined maximum values selected by clinicians. In response to the acoustic feedback, patients were instructed to correct their position, time at which the acoustic feedback ceased. This strategy had been previously proved with promising results in children with spastic diplegia [9].

#### Motivation and inclusion of challenges

In the field of physical rehabilitation, especially in childhood disability, the F-words (Function, Family, Fitness, Fun, Friends and Future) defined by Dr. Rosenbaum [21] become really important. It is essential to maintain a high patients’ motivation because this concept could affect treatment outcomes [22]. To address this issue, we introduced challenges with goals in each session of the robot-based therapy with the aim of having a more engaged user. An example of that is a classification board where the children could follow their progression along the different sessions, and they were rewarded when the goals of each session were correctly fulfilled.

Moreover, with the same objective of enhancing motivation, the data collected with the robotic platform was explained to the patients through graphics so they could feel part of the team, and their interest in the treatment increased.

The patient’s motivation was subjectively measured in each session through a scale from 0 to 10 points, with 0 being no motivation and 10 being maximum motivation.

### Metrics

In order to objectively measure the patient’s evolution and due to the lack of homogeneity among children with CP, authors decided to evaluate the progression of the therapy by comparing each patient to himself, instead of maintaining a control group. We carried out some analyzes and evaluation metrics in different occasions of the study (Table 4): *during* the use of the robot, before the treatment begins (*pre*), in the *middle* and after the whole sessions (*post*).

**Table 4.**
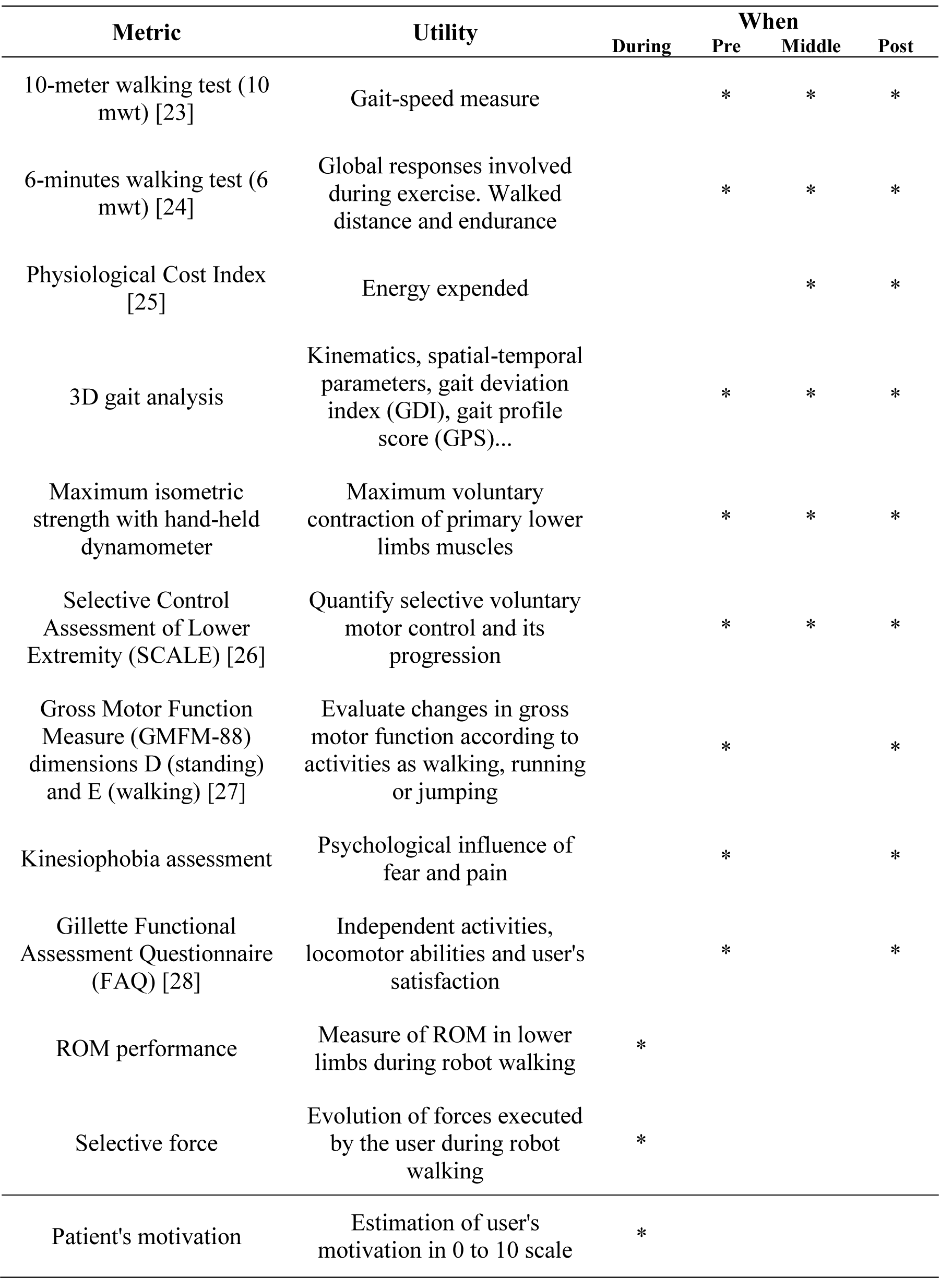
**Evaluation metrics and moment of application**

The 10mwt was assessed for two situations: normal comfortable walking speed and maximum walking speed. Three trials were collected for each situation and subsequently, the average of the three trials was calculated.

Regarding the 6mwt, it was performed indoors along a flat corridor, where the walking course had a 30 m length and was marked every 3 m. The turnaround points were marked with cones. The patients received information about remaining time every minute, but they were not encouraged during the exercise [29]. The heart rate was also measured for each patient in two situations: resting and just after finishing the test. This parameter gives the possibility of calculating the Physiological Cost Index (PCI) after the exercise, which is used to quantify the energy expended by the patients during the exercise and their progression, [25].

3D gait analysis was recorded at 200 Hz using a motion capture system Smart-DX (BTS Bioengineering, Italy). In order to obtain gait kinetics, a set of reflective markers were placed over the skin on discrete anatomical sites according to the Helen Hayes Model [30]. Subjects walked barefoot at a self-selected speed.

Maximum isometric strength was measured in kgf with a hand-held dynamometer microFET2 (Hoggan Scientific LLC, USA). Three records were taken and averaged for each movement bilaterally (dorsiflexion, plantarflexion, knee flexion-extension, hip flexion-extension, abduction and adduction).

The particularity of the SCALE assessment [26] was that it was evaluated by the same physiotherapist bilaterally on three occasions (pre, middle and post), with the aim of reducing the subjective error.

The changes of GMFM-88 [27] were collected for the 88 items, but the comparison analyzes were implemented only for dimensions D (standing) and E (walking).

The kinesiophobia assessment consisted of a test composed of 10 questions of 1 to 4 points each. The responses were given by the patients without parents influence.

Two FAQ questionnaires were requested: one as initial questionnaire at the beginning and the other as follow up at the end of the treatment. These surveys presented several questions for parents and others referred to children.

During the whole treatment, ROM performance and force interactions were measured for each session in order to evaluate if the patient was prepared to jump to the next stage with more difficult parameters and level of assistance.

Finally, the users’ motivation was subjectively evaluated by the practitioner from 0 to 10 points for each session with the robot.

### Patients

Four children diagnosed with spastic CP affecting muscle strength and motor control of lower limbs (two male, two female, weight 44.75±6.29 kg, height 1.56±0.29 m and age 14.50±2.38 years-old) were selected to be participants for testing the robotic training proposal (P1, P2, P3 and P4 in Table 5). The inclusion criteria for patients’ recruitment followed: i) children aged 11 to 18 years suffering from spastic diplegia; ii) Gross Motor Function Classification System (GMFCS [1]) levels I to IV; iii) maximum weight 75 kg; iv) anthropometric measures of lower limbs according to the exoskeleton of CPWalker; v) capable of understanding the proposed exercises; and vi) able to signal pain or discomfort. The exclusion criteria was: i) patients who experimented concomitant treatments 3-months prior study (e.g. orthopedic surgery or botulinum toxin); ii) children with muscle-skeletal deformities or unhealed skin lesions in the lower limbs that could prevent the use of the exoskeleton; iii) patients with critical alterations of motor control as dystonia, choreoathetosis or ataxia; iv) aggressive or self-harming behaviors; and v) severe cognitive impairment. The study was carried out at “Hospital Infantil Universitario Niño Jesús”, (Spain). The Local Ethical Committee of this hospital gave approval to the study (R-0032/12) and warranted its accordance with the Declaration of Helsinki. All participants and families were informed, and parental consents were obtained prior to participation.

**Table 5.** **Patients’ description. Two females (F) and two males (M) with spastic diplegia were selected. No medication 3-months prior the study was taken by the patients. The type of walking support without the aid of the robot is indicated for a distance of 50m and 500m: crutches-CT, wheelchair-WC, posterior walker-PW and cane**.

| Patient | Age | GMFCS | Weight (kg) | Height (m) | Walking support |  |
| --- | --- | --- | --- | --- | --- | --- |
|  |  |  |  |  | 50m | 500m |
| P1 (F) | 12 | III | 40 | 1.56 | CT | WC |
| P2 (F) | 16 | II | 42 | 1.60 | Cane | Cane |
| P3 (M) | 17 | III | 54 | 1.53 | PW | WC |
| P4 (M) | 13 | II | 43 | 1.55 | - | - |

### Results

Due to personal reasons unrelated to the study, three of four patients (P1, P2 and P4) completed 15 of a possible 16 total sessions. Concretely, P1 lost the session number 8, P2 lost the number 7 and P4 the number 11. The rest of training was completed successfully.

The modifications of parameters (ROM, PBWS and gait velocity) proposed in Figure 3 and Figure 4, were fulfilled by all children without problems. The progressions of the levels of assistance provided by Table 3, which were tailored for each patient between sessions 9 to 16 (*power training* with AAN strategies), are represented in Figure 5, where the maximum reached level was level 5 by P1 in the last two sessions.

**Fig. 5.**
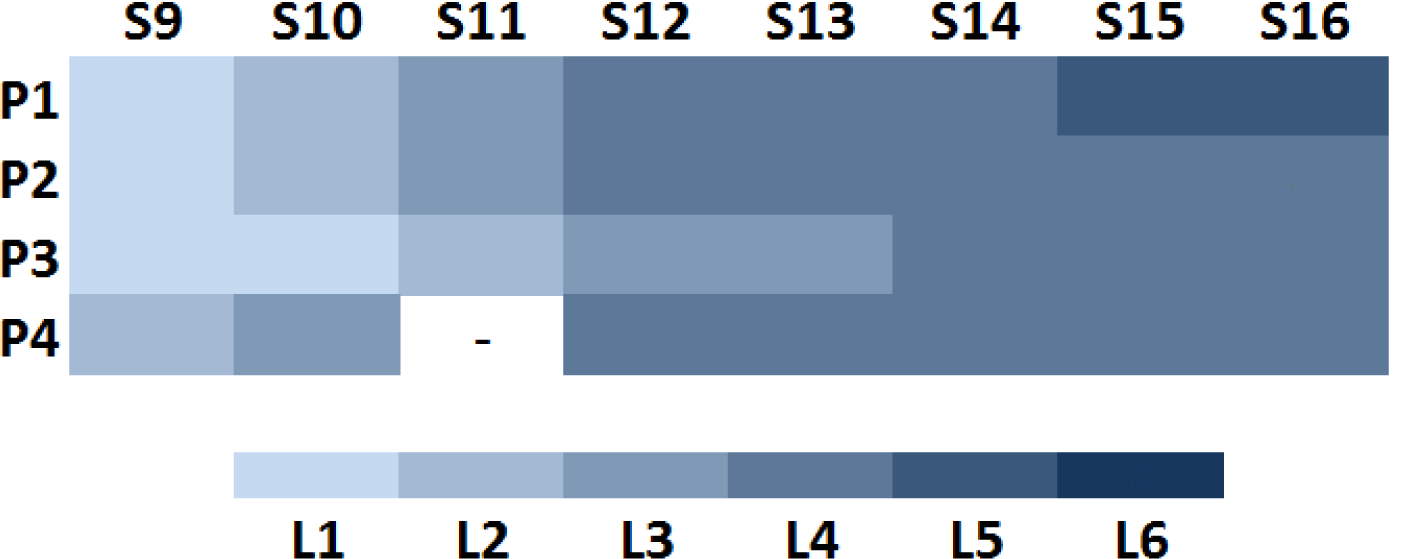
**Levels of assistance (L1 to L6) depending on the patient (P1 to P4) and the AAN session (S9 to S16). The patients could jump to the next level if they achieved at least the 85% of the pattern desired for each session. The level for P4 in S11 is not represented because P4 lost this session**.

### Gait speed, endurance and global responses

All patients improved the outcomes in D and E dimensions of the GMFM-88 scale [27] (Figure 6 a). Results, comparing pre and post studies, show an improvement in this scale of 36.60% for P1, 1.87% for P2, 20.81% for P3 and 16.23% for P4 (Figure 6 a). The SCALE assessment also showed better results at the end of the robot-based treatment (Figure 6 b), [26]. In this case, although the value for the left leg in P1 was kept same as at the beginning (Figure 6 b, red bars for P1), the rest of measures were increased or maintained as maximum (SCALE equal to 10 points).

**Fig. 6.**
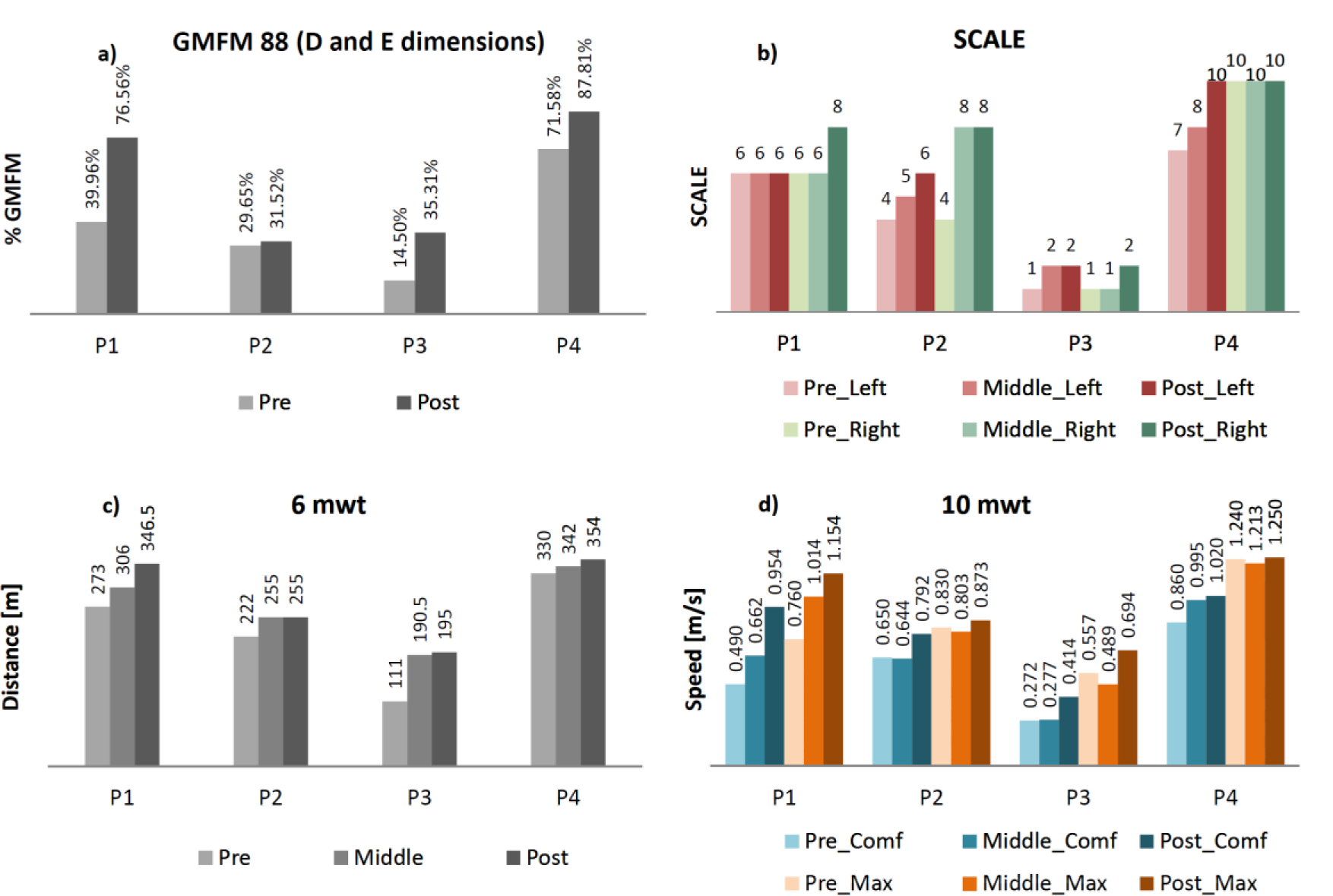
**(a) Results of GMFM-88 (D and E dimensions), (b) SCALE, (c) 6mwt, and (d) 10mwt in pre, middle and post analysis for all the patients (P1 to P4). The SCALE was measured bilaterally (left and right). The 10mwt was performed in two situations: comfortable speed (Comf) and maximum speed (Max)**.

Finally, both the walked distance in the 6mwt and the walking speed in the 10mwt increased after the training period (Figure 6 c and d respectively). The two situations evaluated for the 10mwt are represented: a comfortable speed for each child (blue bars in Figure 6 d) and the same exercise at maximum speed (orange bars in Figure 6 d). More concretely, the percentage of progressions comparing post and pre-analysis in these metrics, were: P1 (6mwt: 26.92%; 10mwt_comf_: 94.69%; 10mwt_max_: 51.84%); P2 (6mwt: 14.86%; 10mwt_comf_: 21.85%; 10mwt_max_: 5.18%); P3 (6mwt: 75.68%; 10mwt_comf_: 52.21%; 10mwt_max_: 24.60%) and P4 (6mwt: 7.27%; 10mwt_comf_: 18.60%; 10mwt_max_: 0.81%).

The Physiological Cost Index was evaluated comparing middle and post assessments during the 6mwt. All patients reduced the PCI: P1 obtained 0.75 beats/m (middle) and 0.55 beats/m (post); P2 0.89 beats/m (middle) and 0.80 beats/m (post); P3 1.57 beats/m (middle) and 1.26 beats/m (post); and P4 0.33 beats/m (middle) and 0.03 beats/m (post).

### Strength progression

In order to quantify the patient’s maximum strength performing on defined and individual movements without the robot, three measures were taken for each required motion. According to that, Figure 7 represents the average values (in kgf) of the individualized movements recorded in pre, medium and post analyzes. In general, results from Figure 7 show that the purple line (post assessment) covers the light-pink line (pre measures) for all the patients. In some cases, it even covers the dark-pink line (measured just after finishing the first 8 strength training sessions). This means that higher values of strength were reached by the children after the robot-based treatment. Concretely, the general improvements (including all the required movements) per patient in maximum isometric strength measure, comparing pre and post averages, were: P1: 129.77±58.71%; P2: 61.39±58.55%; P3: 70.54±83.68% and P4: 34.41±30.41%.

**Fig. 7.**
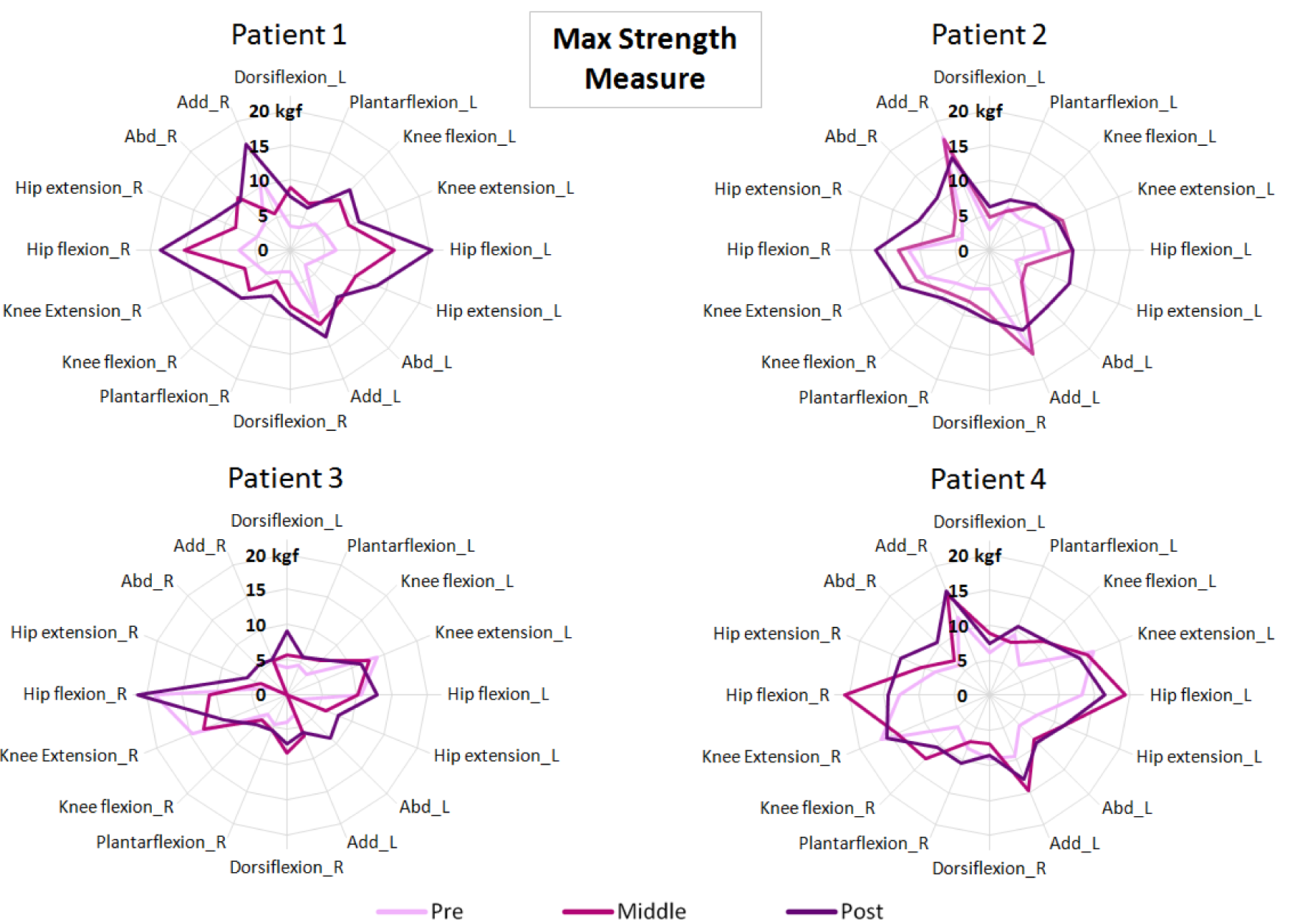
**Maximum strength measures recorded for all the patients in pre (the light-pink line), middle (the dark-pink line) and post analysis (the purple line). Both legs were evaluated, right (R) and left (L)**.

### Kinematics and spatiotemporal variability

The 3D kinematic analysis provided outcomes focused on gait improvements respect to normality. The GPS and GDI (Figure 8) are accepted indexes that represent how close the patient’s gait is to the desired gait. Related to these metrics and comparing pre and post analyzes, all the patients obtained better values for both sides (left and right) after the robot-based treatment. Nevertheless, they were not clinically significant, except for the right side of P1 (around 10 points in GDI). It is important to highlight that we believe that the post results in P4 could be affected by personal circumstances non-related to the study that occurred the day of the test.

**Fig. 8.**
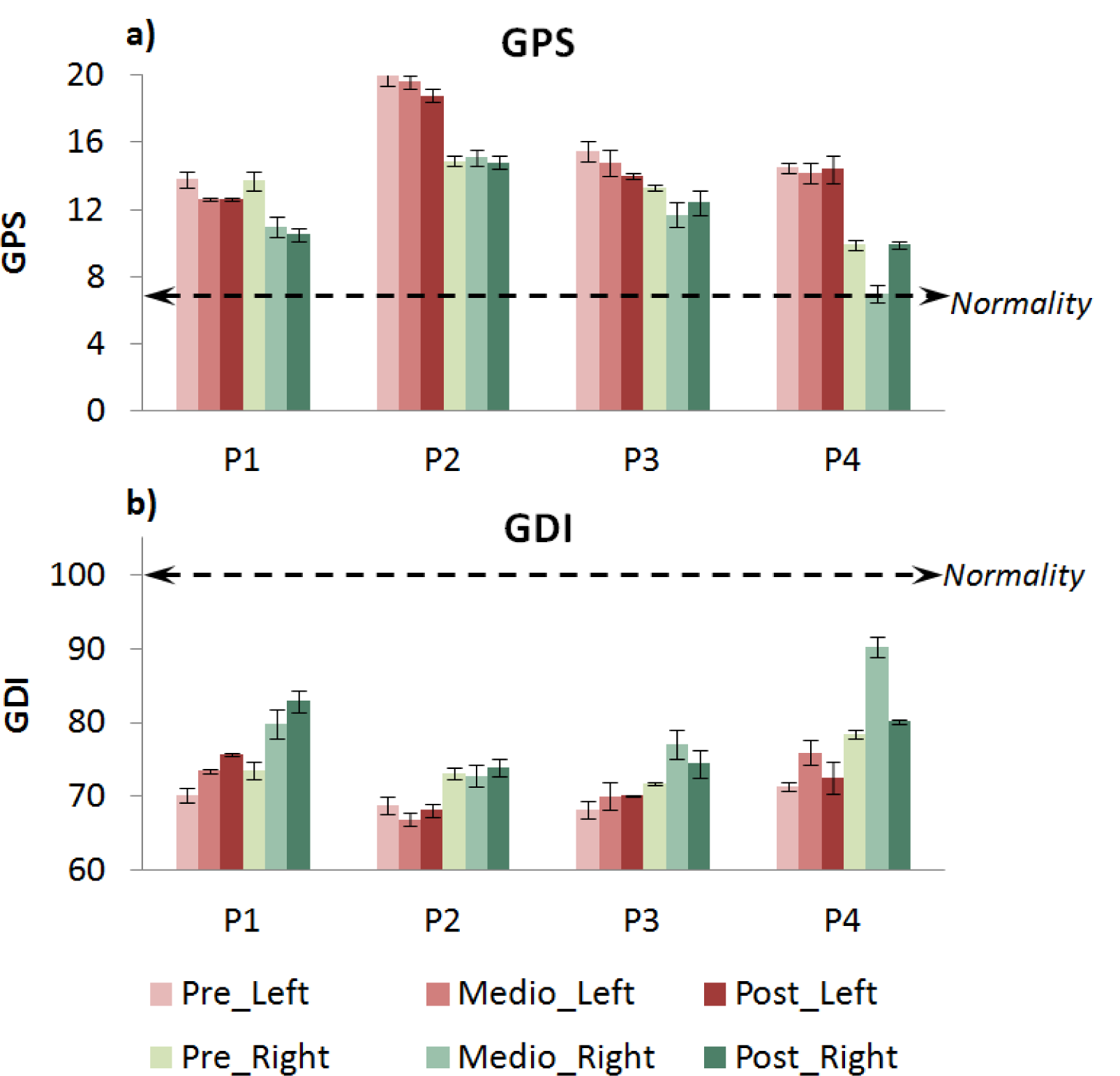
**(a) GPS and (b) GDI for pre, middle and post kinematic analyzes of patients P1 to P4. The results represent means ± standard error bilaterally (left in red bars and right in green bars). Normality in GPS considers values lower than 7 points (doted-black line in (a)), and normality in GDI comprehends values higher than 100 points (doted-black line in (b))**.

Table 6 shows the values of Figure 8 in detail and also includes some of the spatial-temporal parameters recorded during the studies. The average improvement percentages (four patients) in spatiotemporal parameters were: 21.46±33.79% for mean velocity, 2.84±13.96% for cadence and 17.95±20.45% for step length.

### ROM performance

Although all the patients succeeded the changes in the parameters of ROM, velocity and PBWS for the different sessions, the progression in AAN levels was individualized for each subject (Figure 5). Thereby, P1 was the most advanced, achieving level 5 as the maximum. An example of ROM performance difference between trajectory tracking and AAN strategy is represented by Figure 9, which shows the data collected for P1 in session 3 (Figure 9 a) and session 16 (Figure 9 b). The last one was recorded within level 5 (low impedance at hips and medium impedance at knees).

**Fig. 9.**
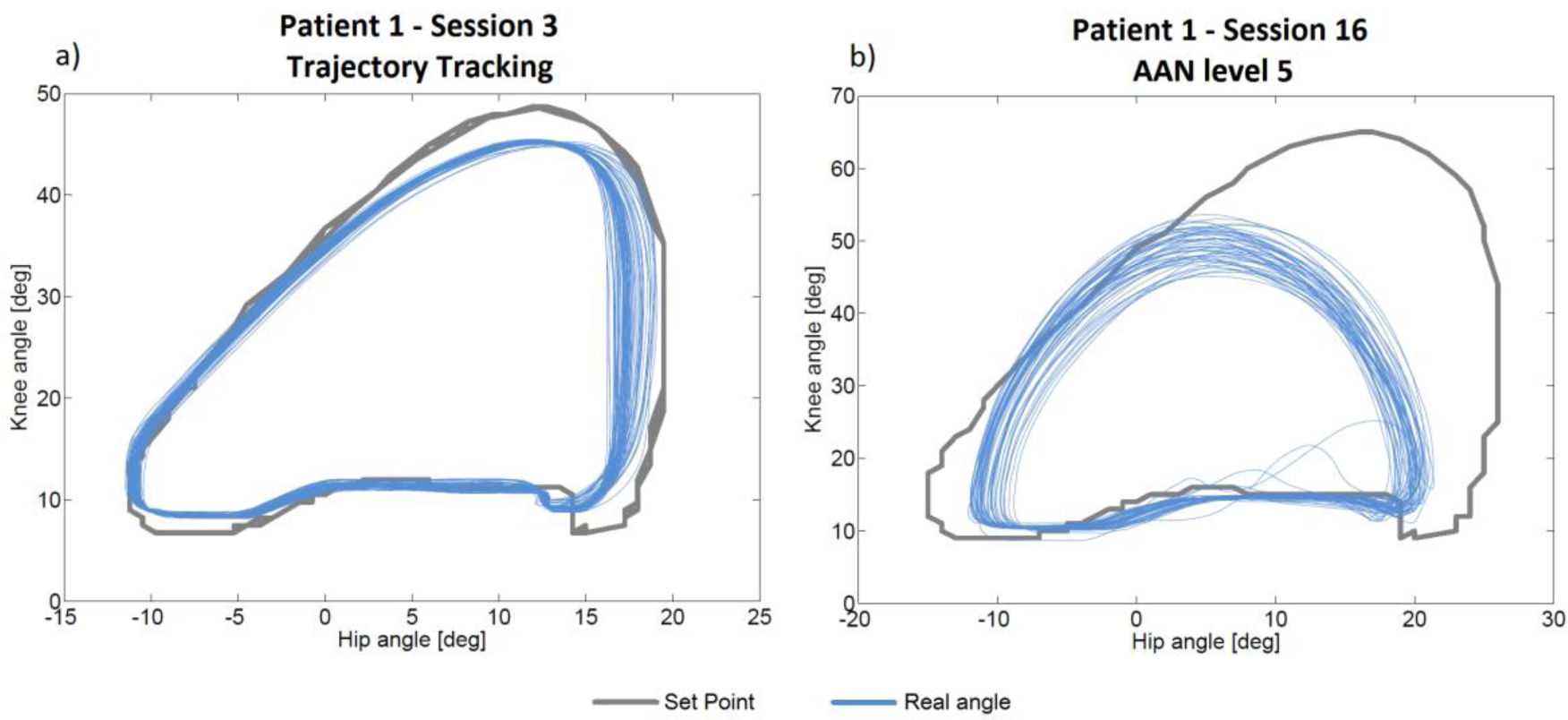
**ROM performance measured walking some steps with CPWalker in two situations: a) Session 3 for patient 1 (*first phase* of the training with trajectory tracking) and b) Session 16 for patient 1 (*second phase* of the training through AAN strategies with level 5 of assistance (LI at hips and MI at knees))**.

### Qualitative variables

The motivation was subjectively evaluated by the practitioner who was with the children during the whole period of the study. The averaged motivation values on a scale from 0 to 10 points were: 9.4 for P1, 8.6 for P2, 9.44 for P3 and 8.87 for P4. Moreover, three of four patients decreased the kinesiophobia score after the 16 sessions. Parents and patients filled a FAQ questionnaire at the beginning and a follow-up at the end of the treatment. Regarding parent questionnaires, results show that all of them thought that the strength and mobility were better at the end of the study thanks to the robot-based therapy. Meanwhile half of them also included the endurance as an improved variable due to the robot. 100% of parents felt satisfied towards the results of robotic therapy with CPWalker, and they ensured that they would like to do it again. The patients’ opinion was very similar. They were also satisfied and, in general, they described the treatment as: “really fun”, “the robot makes you feel light and independent” and “safe”.

## Discussion and Conclusion

The main aim of this research was to provide a first approach to the implementation of a novel and defined robotic rehabilitation method that could cover the most important clinical aspects of the ICF-CY framework. This proposal was tested with four pediatric patients with CP, which provided us some preliminary outcomes to assess it. Although the patients’ progression was evaluated without a control group, we do not consider it as a relevant limitation of the study, since the wide variety of differences among each child with CP makes interesting and even necessary to expose the improvements by comparing each patient with himself.

According to the results, the greatest benefits due to the robot-based treatment corresponded to P1 and P3, who were the most affected levels of GMFCS (III in both cases). In general, the higher values of gait speed and improved values of global responses achieved by all the children in several tests, may be in benefit of the patients’ social mobility. Visual inspection of the graphics show that changes appeared after a small number of sessions (middle tests) and they were commonly increased or maintained until the post studies.

The most challenging part was the *second phase* of the training, which allowed the possibility of adapting the level of assistance depending on the patient’s progression. Thereby, any subject could achieve the last level (level 6), and although the action of reaching level 4 took P3 longer than the rest of children, this patient could pass through it in the last three sessions.

It is interesting to highlight that the outcomes from isometric strength measure showed important peaks of improvement, especially for hip and knee flexion-extension, which was targeted with the CPWalker robotic platform. These higher values were observed from the middle to the post analysis. The results of the present study are difficult to compare with the scientific literature due to the lack of studies using exoskeletons for gait resistance training. However, we can establish that our results are in line with previous studies assessing conventional resistance strength training in CP [31].

In relation to 3D-kinematic analysis, as we said before, P4 suffered a non-grata personal situation the day of the post study. We believe this affected the results of this metric. Nevertheless, the whole population improved the values for spatiotemporal parameters, GDI and GPS, although some of these improvements were not clinically significant. It is important to highlight that not only the kinematics was improved, also the physiological expenditure in walking activities decreased, as illustrated by the reduction of the PCI in all patients, which means that better gait performance implies lower energy cost.

Finally, the motivation scale for the patients and the parents’ satisfaction with the robot-based treatment was very high in most aspects.

The greatest achievements of this proposal come from the possibility of exercising different gait functions in an orderly way, individualized per joint and at the same time than over-ground walking. The proposed protocol could be applied to any current robotic device for gait rehabilitation making minimal changes on it: e.g. in treadmill pediatric platforms as Lokomat [14], despite the impossibility of over-ground walking, it already has different controllers, whose operation modes are close to the levels of impedance of CPWalker, ensuring the progression of the therapy into the sessions. Regarding working postural control in parallel with lower limbs training, which we considered one of the key factors of the study, it may be solved through other solutions if the selected robotic device does not have a similar strategy as described with CPWalker, but it is crucial to guarantee its compliance to get the best results of the treatment [12].

The principal limitation of this research is that we followed up only in short term, so further research with a higher population size is needed to evaluate if the improvements will be kept over time. Moreover, although other studies propose interventions on 3 non-consecutive days per week [31], the patients of the present study performed the robotic exercises during 2 non-consecutive sessions per week. This enabled them to continue their conventional therapies in parallel to the robot-based rehabilitation. The conventional therapies had been attended on a regular basis for years, so authors considered to not abolish them for ethical reasons. The conventional therapies of the patients were performed 2 days a week and consisted of exercises on balance and strength focusing on quadriceps and abs. Although the patients were doing conventional and robotic therapy in parallel, authors consider that the improvements achieved in this proposal are exclusively associated to the use of the CPWalker, since the patients got non-robotic therapies for years with no significant improvements. In conclusion, the method implemented with CPWalker is complementary to the common therapies, providing new possibilities to the clinical practice through robotic rehabilitation and also reaching better outcomes than conventional therapy alone.

We are currently working on new future lines in which we plan to include electromyography during the use of the robotic trainer in order to assess more objectively the patients’ progressions. Electromyography data will serve to evaluate the therapy outcomes in terms of reorganization of the neural structures, which will be performed by analyzing the raw neural drive to muscles and the muscle synergies.

In a nutshell, this manuscript contributed with a defined robotic treatment that could be implemented in most of the existing rehabilitation robotic devices for lower limbs, and which we evaluated positively in four patients with CP using CPWalker.

**Table 6.**
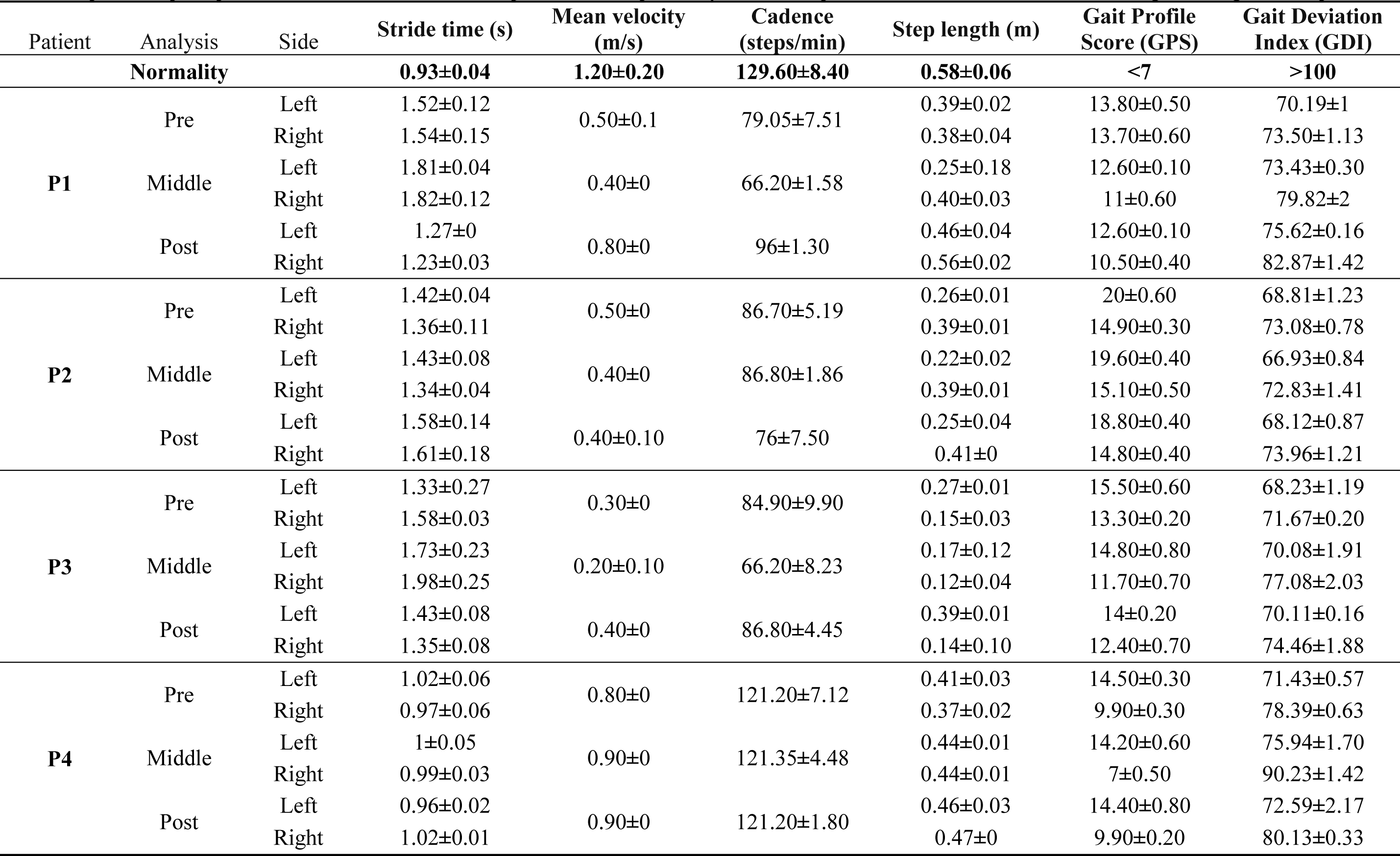
**Spatial-temporal parameters, GPS and GDI values for pre, middle and post analyzes in all the patients. The results were calculated taking an average of 40 steps**.

| Patient | Analysis | Side | Stride time (s) | Mean velocity (m/s) | Cadence (steps/min) | Step length (m) | Gait Profile Score (GPS) | Gait Deviation Index (GDI) |
| --- | --- | --- | --- | --- | --- | --- | --- | --- |
| **Normality** |  |  | **0.93±0.04** | **1.20±0.20** | **129.60±8.40** | **0.58±0.06** | **<7** | **>100** |
| **P1** | Pre | Left | 1.52±0.12 | 0.50±0.1 | 79.05±7.51 | 0.39±0.02 | 13.80±0.50 | 70.19±1 |
|  |  | Right | 1.54±0.15 |  |  | 0.38±0.04 | 13.70±0.60 | 73.50±1.13 |
|  | Middle | Left | 1.81±0.04 | 0.40±0 | 66.20±1.58 | 0.25±0.18 | 12.60±0.10 | 73.43±0.30 |
|  |  | Right | 1.82±0.12 |  |  | 0.40±0.03 | 11±0.60 | 79.82±2 |
|  | Post | Left | 1.27±0 | 0.80±0 | 96±1.30 | 0.46±0.04 | 12.60±0.10 | 75.62±0.16 |
|  |  | Right | 1.23±0.03 |  |  | 0.56±0.02 | 10.50±0.40 | 82.87±1.42 |
| **P2** | Pre | Left | 1.42±0.04 | 0.50±0 | 86.70±5.19 | 0.26±0.01 | 20±0.60 | 68.81±1.23 |
|  |  | Right | 1.36±0.11 |  |  | 0.39±0.01 | 14.90±0.30 | 73.08±0.78 |
|  | Middle | Left | 1.43±0.08 | 0.40±0 | 86.80±1.86 | 0.22±0.02 | 19.60±0.40 | 66.93±0.84 |
|  |  | Right | 1.34±0.04 |  |  | 0.39±0.01 | 15.10±0.50 | 72.83±1.41 |
|  | Post | Left | 1.58±0.14 | 0.40±0.10 | 76±7.50 | 0.25±0.04 | 18.80±0.40 | 68.12±0.87 |
|  |  | Right | 1.61±0.18 |  |  | 0.41±0 | 14.80±0.40 | 73.96±1.21 |
| **P3** | Pre | Left | 1.33±0.27 | 0.30±0 | 84.90±9.90 | 0.27±0.01 | 15.50±0.60 | 68.23±1.19 |
|  |  | Right | 1.58±0.03 |  |  | 0.15±0.03 | 13.30±0.20 | 71.67±0.20 |
|  | Middle | Left | 1.73±0.23 | 0.20±0.10 | 66.20±8.23 | 0.17±0.12 | 14.80±0.80 | 70.08±1.91 |
|  |  | Right | 1.98±0.25 |  |  | 0.12±0.04 | 11.70±0.70 | 77.08±2.03 |
|  | Post | Left | 1.43±0.08 | 0.40±0 | 86.80±4.45 | 0.39±0.01 | 14±0.20 | 70.11±0.16 |
|  |  | Right | 1.35±0.08 |  |  | 0.14±0.10 | 12.40±0.70 | 74.46±1.88 |
| **P4** | Pre | Left | 1.02±0.06 | 0.80±0 | 121.20±7.12 | 0.41±0.03 | 14.50±0.30 | 71.43±0.57 |
|  |  | Right | 0.97±0.06 |  |  | 0.37±0.02 | 9.90±0.30 | 78.39±0.63 |
|  | Middle | Left | 1±0.05 | 0.90±0 | 121.35±4.48 | 0.44±0.01 | 14.20±0.60 | 75.94±1.70 |
|  |  | Right | 0.99±0.03 |  |  | 0.44±0.01 | 7±0.50 | 90.23±1.42 |
|  | Post | Left | 0.96±0.02 | 0.90±0 | 121.20±1.80 | 0.46±0.03 | 14.40±0.80 | 72.59±2.17 |
|  |  | Right | 1.02±0.01 |  |  | 0.47±0 | 9.90±0.20 | 80.13±0.33 |

## Declarations

## List of abbreviations

CP: Cerebral Palsy
CNS: central nervous system
ICF-CY: International Classification of Functioning, Disability and Health framework, Children and Youth version
PBWS: partial body weight support
AAN: assist as needed
ROM: range of motion
NSCA: National Strength and Conditioning Association
IMUs: inertial measurement units
10mwt: 10 meters walking test
6mwt: 6 minutes walking test
GDI: gait deviation index
GPS: gait profile score
SCALE: Selective Control Assessment of Lower Extremity
GMFM: Gross Motor Function Measure
FAQ: Functional Assessment Questionnaire
PCI: Physiological cost index
GMFCS: Gross Motor Function Classification System.

## Ethics approval and consent to participate

The Local Ethical Committee of the “Hospital Infantil Universitario Niño Jesús” gave approval to the study and warranted its accordance with the Declaration of Helsinki. The study was carried out with the number R-0032/12 from Local Ethical Committee of the Hospital, and was publicly registered with the number ISRCTN18254257 as clinical trial. All patients and families were informed beforehand and provided consent through parents to participate.

## Consent for publication

Consent for publication has been given by parents or legal guardians of involved patients.

## Availability of data and materials

The data supporting the conclusions of this article are included within the article and its additional files.

## Authors’ contribution

CB collaborated in the design of the training program, developed the control algorithm of the robotic device, processed the clinical data and drafted the manuscript.

TM collaborated in the design of training program proposal and reviewed the manuscript.

BM collaborated in the clinical processing data derived from the study and helped with the recording of clinical metrics.

OR was involved in the development of the hardware of the robotic device.

APS participated in patients’ recruitment and collaborated in the recording of clinical metrics.

SLL participated in patients’ recruitment and collaborated in the review and critique of the manuscript.

IM aided in the patient’s recruitment and reviewed and criticized the manuscript.

ER collaborated in all facets of the contribution.

ALL AUTHORS read and approved the final manuscript

## Acknowledgements

Authors would like to thank Biomechanical Institute of Valencia and other colleges from Spanish National Research Council and Niño Jesús Hospital for their collaboration in other aspects and achievements carried out with CPWalker robotic platform. Authors appreciate Dr. Gaebler’s advice for the development of the robotic program proposal.

We would like to thank *Made for Movement* company for supporting us with *NF-Walker* device.

We also acknowledge the effort and contribution from all testing children and their families.

We acknowledge support of the publication fee by the CSIC Open Access Publication Support Initiative through its Unit of Information Resources for Research (URICI).

## References

[1] Palisano R, Rosenbaum P, Walter S, Russell D, Wood E, Galuppi B. Gross Motor Function Classification System. Dev Med Child Neurol 1997;39:214–23.

[2] Dietz V. Clinical Aspects for the Application of Robotics in Locomotor Neurorehabilitation. In: Reinkensmeyer DJ, Dietz V, editors. Neurorehabilitation Technol., Springer; 2016, p. 209–21. doi:10.1007/978-3-319-28603-7_11.

[3] Meyer-Heim A, van Hedel HJ a. Robot-assisted and computer-enhanced therapies for children with cerebral palsy: current state and clinical implementation. Semin Pediatr Neurol 2013;20:139–45. doi:10.1016/j.spen.2013.06.006.

[4] Bayón C, Raya R, Lara SL, Ramírez Ó, Serrano J, Rocon E. Robotic Therapies for Children with Cerebral Palsy: A Systematic Review. Transl Biomed 2016;7:1–10. doi:10.21767/2172-0479.100044.

[5] Diaz I, Gil J, Sanchez E. Lower-limb Robotic Rehabilitation: literature review and challenges. J Robot 2011.

[6] van Asseldonk EHF, van der Kooij H. Robot-Aided Gait Training with LOPES. In: Dietz V, editor. Neurorehabilitation Technol., London: Springer London; 2012, p. 379–96. doi:10.1007/978-1-4471-2277-7_21.

[7] Bayón C, Ramírez O, Serrano JI, Castillo MD Del, Pérez-Somarriba A, Belda-Lois JM, et al. Development and evaluation of a novel robotic platform for gait rehabilitation in patients with Cerebral Palsy: CPWalker. Rob Auton Syst 2017;91:101–14. doi:10.1016/j.robot.2016.12.015.

[8] Bartenbach V, Gort M, Riener R. Concept and Design of a Modular Lower Limb Exoskeleton. 6th IEEE RAS/EMBS Int. Conf. Biomed. Robot. Biomechatronics, 2016, p. 649–54. doi:10.1109/BIOROB.2016.7523699.

[9] Bayón C, Lerma S, Ramírez O, Serrano JI, Del Castillo MD, Raya R, et al. Locomotor training through a novel robotic platform for gait rehabilitation in pediatric population: short report. J Neuroeng Rehabil 2016;13:98. doi:10.1186/s12984-016-0206-x.

[10] Rüdt S, Moos M, Seppey S, Riener R, Marchal-Crespo L. Towards More Efficient Robotic Gait Training: A Novel Controller to Modulate Movement Errors. 6th IEEE RAS/EMBS Int Conf Biomed Robot Biomechatronics 2016:884–9.

[11] Lefmann S, Russo R, Hillier S. The effectiveness of robotic-assisted gait training for paediatric gait disorders: systematic review. J Neuroeng Rehabil 2017;14:1. doi:10.1186/s12984-016-0214-x.

[12] Wallard L, Dietrich G, Kerlirzin Y, Bredin J. Robotic-assisted gait training improves walking abilities in diplegic children with cerebral palsy. Eur J Paediatr Neurol 2017:1–8. doi:10.1016/j.ejpn.2017.01.012.

[13] World Health Organization. International Classification of Functioning, Disability and Health, children and youth version. World Health Organization; 2007.

[14] Aurich-Schuler T, Grob F, Van Hedel HJA, Labruyère R. Can Lokomat therapy with children and adolescents be improved? An adaptive clinical pilot trial comparing Guidance force, Path control, and FreeD. J Neuroeng Rehabil 2017;14:1–14. doi:10.1186/s12984-017-0287-1.

[15] Faigenbaum, Avery D. Kramer, W. Blimkie C. Youth Resistance Training: updated position statement paper from the National Strength and Conditioning Association. J Strength Con Res 2009;23:60–79. doi:10.1519/JSC.0b013e31819df407.

[16] Hussain S, Xie SQ, Liu G. Robot assisted treadmill training?: Mechanisms and training strategies. Med Eng Phys 2011;33:527–33. doi:10.1016/j.medengphy.2010.12.010.

[17] Van Asseldonk EHF, Veneman JF, Ekkelenkamp R, Buurke JH, Van Der Helm FCT, Van Der Kooij H. The effects on kinematics and muscle activity of walking in a robotic gait trainer during zero-force control. IEEE Trans Neural Syst Rehabil Eng 2008;16:360–70. doi:10.1109/TNSRE.2008.925074.

[18] Damiano DL, Abel MF, Dl AD, Mf A. Functional outcomes for strenght training in spastic cerebral palsy. Arch Phys Med Rehabil 1998;79:119–25.

[19] Martín-Lorenzo T, Lerma-Lara S, Bayón C, Ramírez Ó, Rocon E. The CPWalker for Strength Training in Children with Spastic Cerebral Palsy: A training program proposal. Int. Conf. Neurorehabilitation, 2016.

[20] Dewar R, Love S, Johnston LM. Exercise interventions improve postural control in children with cerebral palsy: a systematic review. Dev Med Child Neurol 2014. doi:10.1111/dmcn.12660.

[21] Rosenbaum P, Gorter JW. The “F-words” in childhood disability: I swear this is how we should think! Child Care Health Dev 2012;38:457–63. doi:10.1111/j.1365-2214.2011.01338.x.

[22] Maclean N, Pound P. A critical review of the concept of patient motivation in the literature on physical rehabilitation. Soc Sci Med 2000;50:495–506.

[23] Rehabilitation Measures Database. Timed 10-Meter Walk Test. Timed 10-M Walk Test n.d. www.rehabmeasures.org (accessed August 11, 2016).

[24] Crapo RO, Casaburi R, Coates AL, Enright PL, MacIntyre NR, McKay RT, et al. ATS statement: Guidelines for the six-minute walk test. Am J Respir Crit Care Med 2002;166:111–7. doi:10.1164/rccm.166/1/111.

[25] Butler P, Engelbrecht M, Major RE, Tait JH, Stallard J, Patrick JH. Physiological Cost Index of Walking for Normal Children and Its Use As an Indicator of Physical Handicap. Dev Med Child Neurol 1984;26:607–12. doi:10.1111/j.1469-8749.1984.tb04499.x.

[26] Fowler EG, Staudt LA, Greenberg MB, Oppenheim WL. Selective Control Assessment of the Lower Extremity (SCALE): Development, validation, and interrater reliability of a clinical tool for patients with cerebral palsy. Dev Med Child Neurol 2009;51:607–14. doi:10.1111/j.1469-8749.2008.03186.x.

[27] Russell DJ, Rosenbaum PL, Avery LM, Lane M. Gross motor function measure (GMFM-66 and GMFM-88) user’s manual. Press, Cambridge University; 2002.

[28] Gorton GE, Stout JL, Bagley AM, Bevans K, Novacheck TF, Tucker CA. Gillette Functional Assessment Questionnaire 22-item skill set: Factor and Rasch analyses. Dev Med Child Neurol 2011;53:250–5. doi:10.1111/j.1469-8749.2010.03832.x.

[29] Borg G. Psychophysical bases of perceived exertion. Med Sci Sport Exerc 1982;14:377–81.

[30] Kadaba MP, Ramakrishnan HK, Wootten ME. Measurement of Lower Extremity Kinematics During Level Walking. J Orthop Res 1990;8:383–92.

[31] Park EY, Kim WH. Meta-analysis of the effect of strengthening interventions in individuals with cerebral palsy. Res Dev Disabil 2014;35:239–49. doi:10.1016/j.ridd.2013.10.021.

